# Within-patient evolution of plasmid-mediated antimicrobial resistance

**DOI:** 10.1101/2022.05.31.493991

**Authors:** Javier DelaFuente, Laura Toribio-Celestino, Alfonso Santos-Lopez, Ricardo Leon-Sampedro, Aida Alonso-del Valle, Coloma Costas, Marta Hernandez-Garcia, Lun Cui, Jeronimo Rodriguez-Beltran, David Bikard, Rafael Canton, Alvaro San Millan

**Author notes:** **Correspondence and request for materials** should be addressed to Javier DelaFuente & Alvaro San Millan.

## Abstract

Antibiotic resistance (AMR) in bacteria is a major threat to public health, and one of the key elements in the spread and evolution of AMR in clinical pathogens is the transfer of conjugative plasmids. The drivers of AMR evolution have been extensively studied *in vitro*, but the evolution of plasmid-mediated AMR *in vivo* remains poorly explored. Here, we tracked the evolution of the clinically-relevant plasmid pOXA-48, which confers resistance to the last-resort antibiotics carbapenems, in a large collection of enterobacterial clones isolated from the gut of hospitalised patients. Combining genomic and experimental approaches, we first characterized plasmid diversity and the genotypic and phenotypic effects of multiple plasmid mutations on a common genetic background. Second, using cutting-edge genomic editing in wild-type multidrug resistant enterobacteria, we dissected three cases of within-patient plasmid-mediated AMR evolution. Our results revealed, for the first time, compensatory evolution of plasmid-associated fitness cost, as well as the evolution of enhanced plasmid-mediated AMR, in bacteria evolving within the gut of hospitalised patients. Crucially, we observed that the evolution of plasmid-mediated AMR *in vivo* involves a pivotal trade-off between resistance levels and bacterial fitness. This study highlights the need to develop new evolution-informed approaches to tackle plasmid-mediated AMR dissemination.

## Introduction

Antimicrobial resistance (AMR) in bacteria has emerged as a major global threat to public health^1^. AMR is particularly concerning in clinical settings, where nosocomial infections increase mortality rates among hospitalised patients and raise the costs associated with infection control and management^2^. The gut microbiota of patients is one of the most important hotspots of AMR dissemination and evolution^3^, and a crucial element in this process is the transfer of conjugative plasmids – circular DNA molecules that replicate independently of the bacterial chromosome and can transfer horizontally between bacteria^4^.

Numerous studies in recent years have characterized the evolution of plasmid-mediated AMR, expanding our understanding of how AMR plasmids evolve and persist in bacterial populations. AMR plasmids dramatically enhance bacterial fitness in the presence of antibiotics, and plasmid-mediated resistance can further evolve through changes in plasmid copy number (PCN)^5–7^, mutations or duplications of plasmid-encoded AMR genes^8,9^ or interactions with chromosomal mutations^10^. However, in the absence of antibiotics, plasmid-induced physiological alterations frequently lead to a decrease in bacterial fitness, a phenomenon known as plasmid cost^11,12^. This cost can be mitigated over time through compensatory mutations in the plasmid or chromosome^13–15^. Remarkably, previous studies showed that the costs associated with AMR plasmids mainly arise from the expression of resistance genes^11,16,17^.This insight suggests that bacteria carrying AMR plasmids probably experience a trade-off between fitness in the presence and absence of antibiotics (fitness-resistance trade-off)^18,19^. Despite the importance of these earlier studies, current understanding of the evolution of plasmid-mediated resistance derives almost entirely from highly controlled experiments conducted *in vitro*. The lack of access to suitable bacterial collections of clinical origin, together with the arduousness of performing genetic manipulations with wild-type, multidrug-resistant bacterial isolates, has prevented study of the evolution of plasmid-mediated AMR in clinically relevant real-life scenarios.

Here, we tracked the evolutionary dynamics of plasmid-mediated AMR in the gut microbiota of hospitalised patients. We focused on the widespread pOXA-48-like conjugative plasmids, which constitute one of the most relevant plasmid groups in clinical settings in Europe^20,21^. pOXA-48-like plasmids are found in the order *Enterobacterales*, giving rise to carbapenem-resistant enterobacteria, which were recently reported to be the fastest-growing resistance threat in Europe^22^. We used a previously characterized collection of 224 pOXA-48-carrying enterobacteria isolated over a two-year period from more than 9,000 hospitalised patients at the Ramon y Cajal University Hospital in Madrid, Spain (R-GNOSIS collection, Supp. Fig. 1. For the characterization of pOXA-48-carrying isolates in R-GNOSIS, see^23–25^). We studied multiple pOXA-48 variants carrying distinct mutations and elucidated the evolution of specific associations between pOXA- 48 and wild-type enterobacteria in the gut microbiota of three patients. Our results revealed that the *in vivo* evolution of plasmid-mediated resistance is shaped by interplay between AMR and plasmid-induced fitness costs.

**Figure 1.**
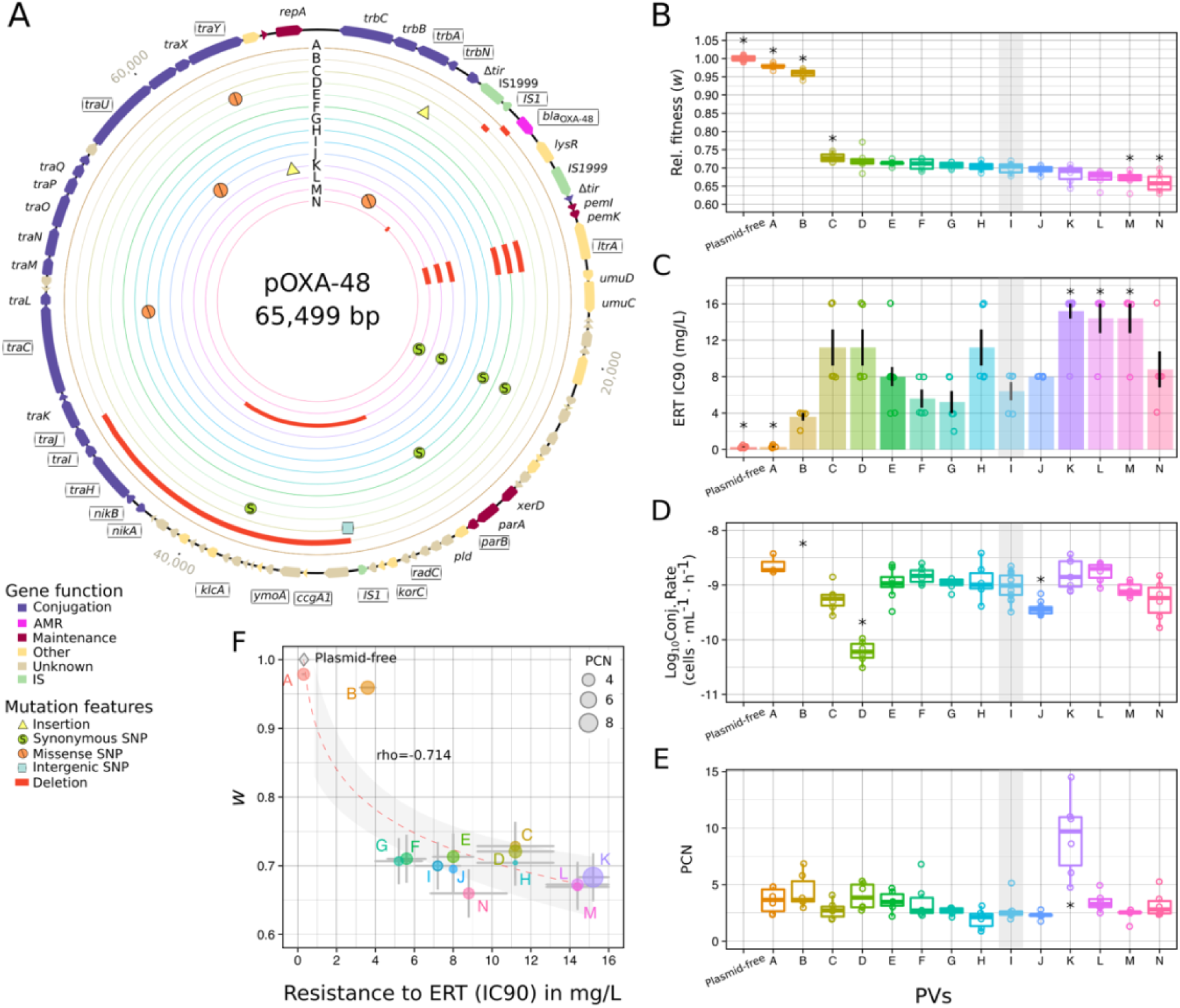
pOXA-48 plasmid variants tested in *E. coli* J53. A) Representation of pOXA-48 variants (PVs) A-N in concentric circles (for variant details see legend). The gene map in the outer circle represents the most common variant (PV-I, used as a reference and highlighted by grey shading in panels B-E). Arrows indicate the reading frames, and colours indicate gene functional classification. Gene names are indicated in the outer circle, and the names of genes showing mutations are represented inside boxes. B) Box plots showing relative fitness (*w*) of pOXA-48-carrying *E. coli* J53 relative to plasmid-free J53. Horizontal lines inside boxes indicate median values, the upper and lower hinges correspond to the 25^th^ and 75^th^ percentiles, and whiskers extend to observations within the 1.5 x the interquartile range (IQR). Individual points show independent replicates (n=6). C) Resistance to ertapenem (ERT) in plasmid-free and plasmid-carrying *E. coli* J53, represented as the 90% inhibitory concentration (IC90) in mg/L. Bars indicate the mean values and dots indicate individual replicates (n=10 for PV-K & PV-E and n=5 for the remaining PVs). Black bars represent the standard error of the mean. D) Conjugation rates of different PVs in *E. coli* J53 (in log_10_ scale, n=14 for PV-I, n=9 for PV-J, n=8 for PV-E & PV-K, n=3 for PV-A, and n=6 for the remaining PVs), represented as boxplots as in B. E) Plasmid copy number of different PVs in *E. coli* J53, represented as boxplots as in B (n=6). Asterisks in panels B-E indicate significance (P<0.05) for the comparison of each PV with PV-I. F) Correlation between relative fitness (*w*) and resistance to ertapenem (mean IC90) in PV-carrying *E. coli* J53. Individual points indicate the mean value, and lines represent the standard error of the mean IC90 and the propagated standard error of the relative fitness. The size of each point is proportional to PCN in J53. The diamond represents the plasmid-free J53 values, which were not included in the correlation. Individual PVs are indicated by letters. The red dashed line indicates the linear regression and the gray-shaded zone covers the 95% confidence interval.

## Results

### Analysis of pOXA-48 plasmid variants

To identify mutations potentially associated with plasmid-mediated AMR evolution, we characterized the genomes of all pOXA-48-like plasmids in the R-GNOSIS collection. Comparison of the full sequences of all 224 pOXA-48-like plasmids, identified a total of 35 plasmid variants (PVs), defined as pOXA-48-like plasmids carrying any SNP or insertion/deletion (indel) compared with the most common variant (PV-I), which is present in ∼67% of the isolates in the collection (Figure 1, Supp. Fig.1, Supp. Table 1 and Methods).

We next studied the phenotypic and genotypic effects of a selection of 14 of the PVs (Figure 1A and see Methods for PV selection criteria). The 14 pOXA-48 variants were introduced into the *Escherichia coli* J53 strain^26^ (K12 derivative), used as a common isogenic bacterial host to specifically dissect plasmid effects. PVs-carrying J53 genomes were resequenced to confirm plasmid presence and the isogenic nature of the transconjugants (Supp. Table 2, see Methods). The following phenotypic and genotypic variables were examined in each transconjugant and in plasmid-free J53: i) bacterial fitness, assessed from growth curves and competition assays; ii) plasmid conjugation rate; iii) antimicrobial resistance; and iv) plasmid copy number (PCN).

The PVs produced a variety of phenotypes in J53 (Figure 1 B-E and Supp. Fig. 2). For example, although the fitness effect of most PVs was similar to that of PV- I (the most common PV), two PVs were associated with a large decrease in cost (Kruskal-Wallis rank-sum test, followed by pairwise comparison Wilcoxon rank-sum exact test with FDR correction P<0.01). One of these variants (PV-A) had a deletion in the carbapenemase gene *bla*_OXA-48_ that abolished plasmid-mediated AMR (Figure 1C). The other one (PV-B) carried two deletions: i) a small deletion affecting the IS1 element upstream of *bla*_OXA-48_ and ii) a ∼13.5 kb deletion including genes involved in conjugation, associated with a conjugation- incompetent phenotype (Figure 1D). PV-D and PV-J both had a lower conjugation rate than PV-I (Figure 1D; Kruskal-Wallis rank-sum test followed by pairwise comparison Wilcoxon rank-sum exact test with FDR correction P<0.001) and carried mutations in conjugation-related genes: a nonsynonymous SNP in *traY* and an insertion in *trbN* (PV-D), and a nonsynonymous SNP in *traU* (PV-J) (Supp. Table 1). PCN was significantly elevated in one PV, PV-K (Figure 1E), which carried a mutation upstream of the gene encoding the replication initiation protein RepA (Kruskal-Wallis rank-sum test, followed by pairwise comparison Wilcoxon rank-sum exact test with FDR correction P=0.022).

Analysis of the effects of AMR, PCN and conjugation rates on plasmid-associated fitness costs in J53 revealed a clear trade-off between AMR and fitness costs (Figure 1F, Spearman’s rank correlation S= 779.7, rho= -0.7136, P= 0.004). PVs conferring AMR were associated with a high fitness cost (27%-34% reduction in relative fitness), whereas the two PVs which conferred low or no AMR imposed only a small fitness cost (<4 % reduction in relative fitness). Neither conjugation nor PCN showed a significant association with plasmid fitness costs (Spearman’s rank correlation, P>0.25, Supp. Fig. 3).

### Screening within-patient pOXA-48 evolution

The above characterization of pOXA-48 variants suggested that PV mutations, and the resulting fitness-resistance trade-off, could contribute to the evolution of pOXA-48-mediated AMR *in vivo*. In the R-GNOSIS project, hospitalised patients colonized with carbapenemase-producing enterobacteria were sampled periodically, generating timelines of isolates that allowed us now to investigate the evolution of pOXA-48-mediated resistance in the gut of these patients. We screened the timeline of bacterial isolates collected from individual patients and selected isolates from the same clonal background but carrying different PVs, which would suggest within-patient plasmid evolution. Among the 121 patients colonized by pOXA-48-carrying enterobacteria, we identified three whose timelines matched these conditions: patients HKH, JWC, and WDV (Figure 2, see methods).

**Figure 2.**
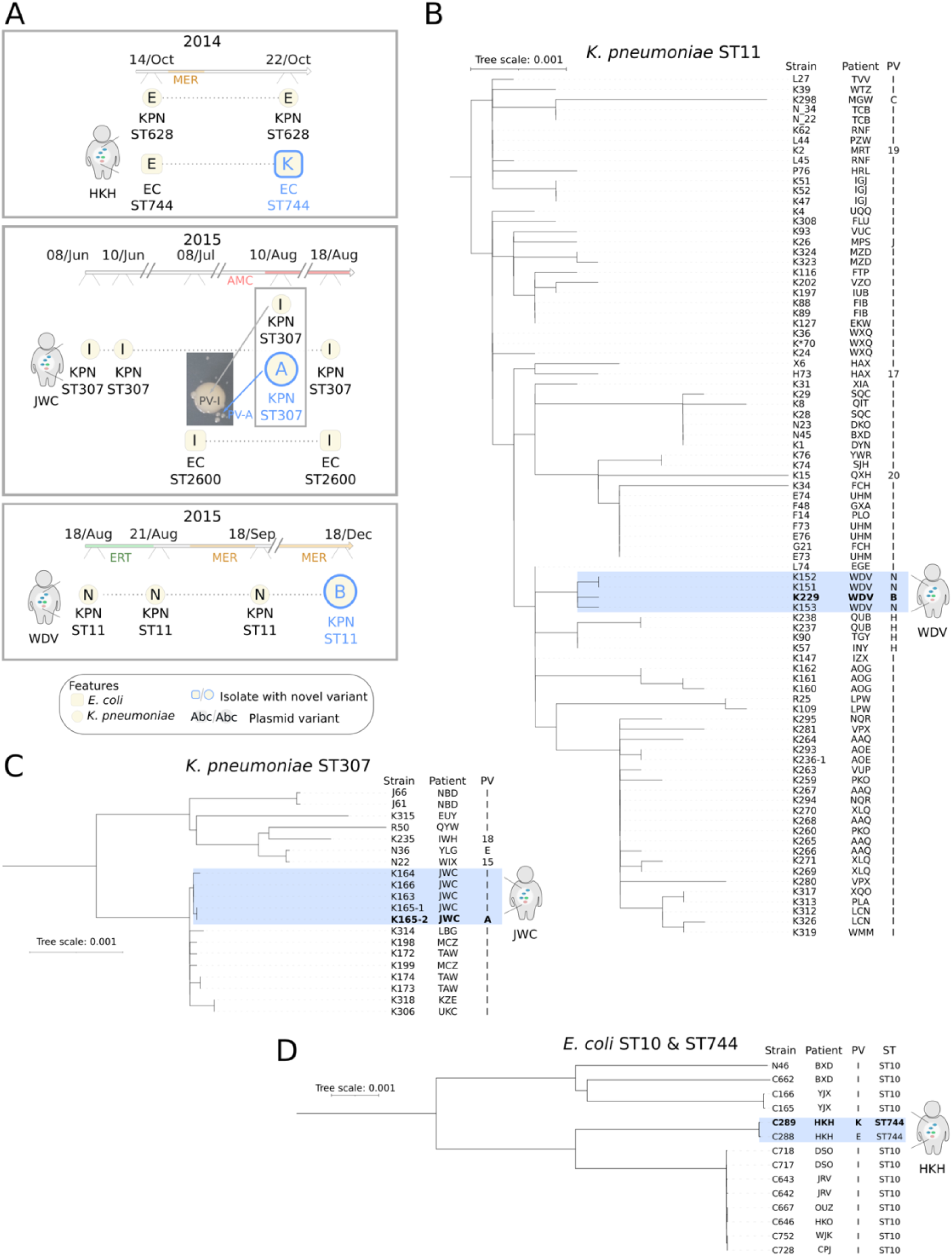
Screening the within-patient evolution of pOXA-48-mediated AMR. A) Timelines of the isolation of pOXA-48-carrying enterobacteria from patients HKH, JWC, and WDV (see legend). Isolate features are detailed in the legend. Swab dates are indicated next to the timeline (day/month). pOXA-48-selecting antibiotic treatments are indicated in the timeline (MER, meropenem; ERT, ertapenem; AMC, amoxicillin + clavulanic acid). Each PVs is indicated by a letter. Species are indicated by letters and symbols (KPN and circle for *K. pneumoniae*; EC and square for *E. coli*). The multilocus sequence-type code is indicated next to the species label. Isolates in which the new PV was detected are indicated by blue type and larger size. The patient JWC timeline reveals the emergence of two *K. pneumoniae* isolates co-isolated on an agar plate supplemented with ertapenem 0.3 mg/L. B-D) Genetic relationship built using core-genome comparisons (midpoint rooted phylogenetic trees) of *K. pneumoniae* ST11 (n=85, B), *K. pneumoniae* ST307 (n=20, C), and *E. coli* ST10 and ST744 (n=12 & n=2 respectively, D) from the collection. Strain designation, patient codes, and PVs are indicated (see Supp table 1). Isolates involved in putative cases of within-patient evolution are highlighted in blue. Bold lettering marks the isolate in which the novel variant was identified. The tree-scale indicates nucleotide substitution per site.

In the genomic analysis of pOXA-48-carrying bacteria from the R-GNOSIS collection, two lines of evidence strongly suggested that plasmid mutations emerged and were subsequently selected in specific clones in the gut microbiota of these three patients. First, comparison of the core genomes of all *K. pneumoniae* and *E. coli* isolates revealed a tight grouping of isolates potentially involved in the potential within-patient evolution events (Figure 2B-D, <5 SNPs between all isolates for each clonal line, Supp. Table 3). This result makes it very unlikely that the observations in patients HKH, JWC, and WDV were the result of independent colonization by clones carrying different PVs. Second, each of the novel PVs originating in these patients was restricted to an individual patient, and was not described in any other isolate in the collection (not even in other clones from the same patient). This result challenges the possibility that these new bacteria-PVs associations were generated by independent conjugation events.

### Evolution of pOXA-48-mediated resistance in vivo

To characterize the within-patient plasmid-mediated AMR evolution, we studied the isolates carrying the novel PVs in each patient (from now on, identified with the patient code followed by an asterisk, Figure 3A). First, we cured the pOXA- 48 PVs from these isolates using an in-house CRISPR-Cas9 system specifically designed to remove plasmids from multidrug-resistant enterobacteria (see methods and Supp. Fig. 4). Then, for each patient, we independently re- introduced both the ancestral PV (the initial PV present in the same clonal line in the same patient) and the novel PVs in these isolates. Crucially, we sequenced the genomes of the wild-type clones by combining long-read and short-read technologies, and resequenced the genomes of all strains after plasmid curing to ensure that no significant mutations occurred during the process (Figure 3A, Supp. Fig. 4 & Supp. Table 3). Once we had introduced the two PVs into each clone, we measured i) plasmid fitness effects, ii) antimicrobial resistance (to all beta-lactam antibiotics used for treatment in these patients), and iii) PCN for every clone (Fig. 3 B-D, Supp. Fig. 4). The results of these analyses are presented in the following sections patient-by-patient in chronological order.

**Figure 3.**
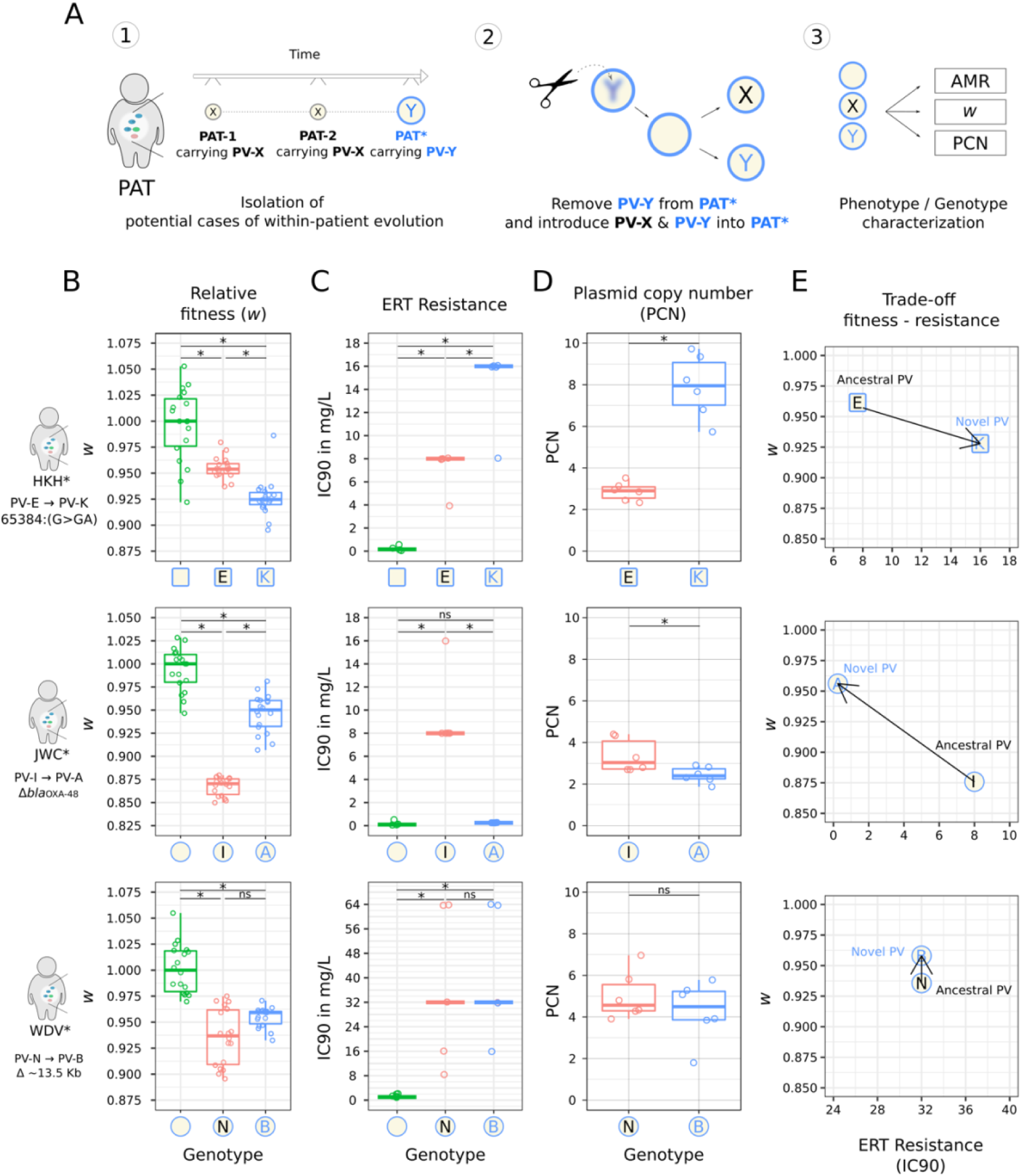
Characterization of the *in vivo* evolution of plasmid-mediated AMR. A) Workflow used to investigate the within-patient evolution of pOXA-48-mediated AMR. B) Relative fitness (*w*) of each plasmid-bacteria combination compared with the plasmid-free strain (see methods). Horizontal lines inside boxes indicate median values, the upper and lower hinges correspond to the 25th and 75th percentiles, and whiskers extend to observations within 1.5 × the IQR. Individual points represent independent replicates (n=18). Asterisks in panels B-D indicate significant differences (P<0.05 in pairwise comparison Wilcoxon rank-sum exact test with FDR correction in B and C, and Padj<0.05 by one-way ANOVA in D); ns, nonsignificant (Padj>0.05). C) Resistance to ertapenem (ERT) measured as IC90 (mg/L) of plasmid-free and plasmid- carrying combinations. Lines indicate median values and individual replicates are indicated by points (n=5). D) Plasmid copy number (PCN) of each PV, represented in boxplots as in B (n=6). E) Schematic representations of the trade-off between antibiotic resistance (median IC90) and relative fitness (median *w*) in the strains under study. Black arrows represent the trade-off and arrowheads indicate the PVs timeline (from ancestral to novel PV).

*Patient HKH. Increased PCN leading to increases in AMR and fitness costs* Four pOXA-48-carrying isolates were recovered from this patient (Figure 2A). Two isolates belonged to a *K. pneumoniae* sequence-type 628 (ST628) clone, and both carried the same plasmid variant (PV-E). The other two isolates belonged to a *Escherichia coli* ST744 clone. The first of the *E. coli* isolates also carried PV-E, while the second (HKH*) was recovered 8 days later and carried a different plasmid variant, PV-K, differing from PV-E by only a single base pair insertion upstream of *repA*. The *in vitro* genotypic characterization in *E. coli* J53 had revealed an association between PV-K and increased PCN (Figure 1E). Analysis of the effects of PV-E and PV-K in HKH* revealed that PV-K was present at a higher PCN (from 3 to 8 copies, one-way ANOVA F=61.42, d.f.=1, Padj<0.001). The high PCN of PV-K in HKH* was associated with increased AMR to ertapenem (Wilcoxon rank-sum test W=2, P=0.02) and meropenem (Supp. Fig. 5 A-B) but decreased fitness in the absence of antibiotics (Wilcoxon rank-sum exact test, W=306, P<0.001, Fig. 2 B-D, Supp. Fig. 5A-B). Patient HKH’s clinical history revealed meropenem treatment before the isolation of HKH*, which carried the high-PCN PV-K.

### Patient JWC. Loss of AMR leading to amelioration of plasmid cost

Over a 10-week period, six pOXA-48-carrying isolates were recovered from patient JWC (Figure 2A). Four of them belonged to a *K. pneumoniae* ST307 clone and the remaining two to an *E. coli* ST2600 clone. Five isolates carried the most common pOXA-48 variant, PV-I, but the *K. pneumoniae* isolate JWC* carried PV- A, which differs from PV-I by a 199 bp deletion starting 163 bp upstream the coding DNA sequence of the *bla*_OXA-48_ gene. As in the J53 analysis, PV-A was associated not only with loss of resistance to ertapenem and amoxicillin- clavulanic acid in JWC* (Fig. 3C and Supp. Fig. 5 C-D), but also with a reduction in plasmid fitness costs compared with PV-I (Wilcoxon rank-sum test W=0, P<0.001 for both phenotypes). PV-A was also associated with a modest but significant decrease in PCN in JWC* (one-way ANOVA F=6.51, d.f.= 1, Padj=0.029, Fig 3 B-D).

In R-GNOSIS, carbapenemase-producing enterobacteria were recovered by selective plating, and it was therefore difficult to understand how the carbapenem- susceptible JWC* isolate was obtained from this patient. To investigate this, we plated the original frozen JWC isolate stocks on agar with and without ertapenem. Antibiotic-containing plates inoculated with the JWC* frozen stock (but not the other isolates) contained large resistant colonies of *K. pneumoniae* ST307/PV-I surrounded by smaller susceptible colonies of isogenic *K. pneumoniae* ST307/PV-A (Figure 2A), an example of the phenomenon of cross-protection by secreted beta-lactamases known as satellitism^27^. Sequencing of the entire genomes of three large and three satellites colonies confirmed that they were isogenic *bla*_OXA-48_+ and *bla*_OXA-48_- variants of the same *K. pneumoniae* ST307 clone. This results strongly suggests that two versions of the *K. pneumoniae* ST307 clone, with the different PVs, coexisted in the patient’s gut at that timepoint. Although PV-A and PV-I carrying bacteria coexisted at the time of JWC* sampling (Figure 2A), 8 days later only the fully resistant PV-I-carrying clone was detected. This shift is probably explained by an AMC treatment that began right before the isolation of JWC*, since OXA-48 confers high-level resistance to AMC.

### Patient WDV. Large plasmid deletion leading to loss of conjugation ability

Four isolates of a *K. pneumoniae* ST11 clone were recovered from this patient over a four-month period. The three initial isolates carried PV-N, but the last isolate (WDV*) carried PV-B, which differed from PV-N by a ∼13.5 kb deletion (Figure 2A and Figure 1A). Compared with PV-I, both PV-N and PV-B carry the same small deletion affecting the IS1 element upstream of *bla*_OXA-48_. The large ∼13.5 kb deletion in PV-B affected multiple genes involved in conjugation, leading to the loss of conjugation ability in J53 (Fig. 1). In the wild-type strain, PV-B was also associated with a conjugation-incompetent phenotype, and it produced a small, marginally significant, decrease in fitness costs in WDV* compared with PV-N (Wilcoxon rank-sum exact test W=101, P=0.054). AMR and PCN were the same in the PV-N-carrying and the PV-B-carrying WDV* isolate (Wilcoxon rank-sum test W=23 & W=14.5, P>0.5). The results from patient WDV thus revealed no clear change in plasmid-associated effects, although the slight difference in fitness costs imposed by PV-N and PV-B could suggests that the large deletion in PV-B might act as a compensatory mutation (Figure 3 B-C and Supp. Fig. 5 E-F).

### A fitness-resistance trade-off shapes within-patient evolution of pOXA-48

In line with our observations in *E. coli* J53, the analysis of the *in vivo* evolution of pOXA-48-mediated AMR in patient gut microbiota indicated that this process is shaped by a fitness-resistance trade-off (Figure 3E). Moreover, clinical metadata from patients strongly suggests that antibiotic treatments direct the rapid, and even bidirectional, navigation of this trade-off.

In patient JWC we detected coexistence of almost isogenic *K. pneumoniae* ST307 populations differing only in the presence of an intact *bla*_OXA-48_ gene in pOXA-48. The presence of the *bla*_OXA-48_ was associated with fitness cost in the absence of antibiotics. The emergence of the *bla*_OXA-48_-lacking PV-A followed a two-month period of no OXA-48-selecting antibiotic treatment, but this variant was rapidly depleted by a cycle of amoxicillin + clavulanic acid. This result not only supports the impact of the fitness-resistance trade-off on the evolution of AMR, but also highlights the importance of clonal diversification in the gut microbiota for this process.

In patient HKH, meropenem treatment triggered the rapid emergence (8 days between isolates) of PV-K, which conferred increased PCN and AMR. Fitness results indicated that PV-K was associated with an increased fitness cost in the absence of antibiotics. Unfortunately, no further samples from this patient were included in the R-GNOSIS collection, and we were therefore unable to investigate the fate of PV-K after the antibiotic treatment ended.

## Discussion

The role of fitness-resistance trade-offs in the evolution of AMR has received considerable attention^18,28^. However, despite plasmids being arguably the most important vehicle for the acquisition of clinically relevant AMR in many key pathogens, there was previously little published evidence of the impact of this trade-off on the evolution of plasmid-mediated AMR in clinically relevant situations^19^. We anticipate that the fitness-resistance trade-off described here will affect the evolution of plasmid-mediated AMR more generally, because AMR gene expression is one of the central sources of plasmid-associated fitness costs^11,16,17^. One naive prediction arising from this result is that, in the absence of antibiotic pressure, natural selection could favour plasmid loss or mutations that inactivate plasmid-encoded resistance genes, reversing AMR evolution. However, this prediction is challenged by at least two lines of evidence. First, we observed that the standing genetic variation in the gut microbiota helps to bypass this fitness-resistance trade-off by supporting the coexistence of subpopulations carrying either low-cost/low-resistance or high-cost/high-resistance PVs. Indeed, because of the sampling and isolation protocol used in R-GNOSIS (isolation of one clone per species and time point), the role of preexisting genetic diversity in AMR evolution is probably vastly underestimated in our analysis. Second, we previously reported that pOXA-48 conjugation is pervasive in hospitalised patients and leads to long-term plasmid carriage in their gut^23^. In that study, we described an *in vivo* pOXA-48 conjugation event in patient HKH involving the same *E. coli* clone in which we have now described subsequent plasmid evolution. The plasmid dynamics described in this patient perfectly exemplify the ability of pOXA-48 to spread rapidly and to evolve in the gut microbiota of hospitalised patients. The high pOXA-48 conjugation rate will feed the microbiota with a constant supply of new plasmid-carrying bacteria, promoting plasmid maintenance in the bacterial community. These results highlight the need to consider AMR ecology and evolution in order to develop more rational strategies to counteract AMR in complex bacterial communities, such as the gut microbiota.

Our study also highlights the need to consider two previously neglected issues in the study of AMR evolution. The first of these is the importance of analysing AMR evolution directly in the wild-type, clinically relevant bacterial strains. Our results showed that although the laboratory *E. coli* strain J53 provides reasonably good qualitative predictions of PVs effects, plasmid-associated fitness costs tend to be much higher than in the wild-type bacteria. This discrepancy could lead to erroneous predictions about the survival of AMR strains in the gut of patients. The second issue is the importance of considering the preexisting genetic diversity in bacterial communities when assessing the potential for AMR evolution. Our results from patient JWC, together with findings from other recent studies^24,29,30^, show that genetic diversity fuels AMR evolution. Most research to date on within- patient evolution of AMR (including this study) has failed properly to screen for community diversity. In future studies, addressing these issues will produce more accurate predictions and help to develop better intervention strategies against AMR in clinical settings.

## Methods

### Bacterial strains, and culture conditions

All experiments were performed in Lennox lysogeny broth (LB) which was -when indicated- supplemented with 15 g/L agar (CONDA, Spain). Mueller Hinton II broth (Oxoid) was used for IC90 determination and results were comparable with those obtained in LB. Amoxicillin+clavulanic acid (Sandoz, Spain), meropenem (Aurovitas, Spain), kanamycin, ertapenem, chloramphenicol, apramycin, streptomycin, sodium azide and carbenicillin (Merck, Spain) were used in this study. Clinical strains used in this study were isolated during the R-GNOSIS project which included 28,089 samples from 9,275 patients in the Hospital Universitario Ramon y Cajal (Madrid, Spain)^23,25^. For this study only pOXA-48- carrying enterobacteria were included. All primers used in the study are described in Supp. Table 4.

### Plasmid construction

pBGC^24^ was used to construct pBGA by exchanging the *cat* gene (chloramphenicol resistance) with *aac(3)-IV* (apramycin resistance) from pMDIAI^31^ by Gibson assembly (New England Biolabs, UK). pLC10-Apra was constructed by exchanging the *aph(3’)-Ia* gene (Kanamycin resistance) with the *aac(3)-IV* gene, by Gibson assembly. Plasmids pLC10-Kan/pLC10-Apra carry the *Streptococcus pyogenes cas9* gene under the control of a Ptet promoter inducible by anhydrotetracycline (derived from pWJ153^32^), cloned on a thermosensitive pSC101 plasmid backbone with a guide RNA under the control of a Ptrc promoter derived from pCas^33^ (Addgene plasmid #62225). Single guide RNA (sgRNA) targeting pOXA-48 *pemK* gene (Fig.1A) was introduced into pLC10-Kan by golden gate assembly^34^ (New England Biolabs, UK). pLC10-Apra was constructed by exchanging the *aph(3’)-Ia* gene (Kanamycin resistance) with the *aac(3)-IV* gene, by Gibson assembly.

### gDNA extraction, short-read (Illumina) and long-read sequencing (MinION)

Genomic DNA was extracted using the Wizard genomic DNA purification kit (Promega). Short-read sequencing data from wild-type strains was obtained from ^23^ (BioProject PRJNA626430). Additionally, *E. coli* J53 transconjugants/transformants and the wild-type strains involved in within-patient pOXA-48 evolution (K163, K165, C288, C289, K153 and K229) were sequenced in the Microbial Genome Sequencing Center (MIGS, USA) using NextSeq 2000 platform (coverage>100x). Long-read sequencing (MinION) was performed in MIGS for the wild-type strains involved in within-patient pOXA-48 (coverage>100x). Sequencing data are available under BioProject PRJNA838107. Short-reads from MiGS were trimmed with Trim Galore v0.6.4 (https://github.com/FelixKrueger/TrimGalore), using a quality threshold of 20 and removing adapters and reads <50 bp. Filtlong v0.2.1 (https://github.com/rrwick/Filtlong) was used for filtering long-reads.

### Assembly and analysis of pOXA-48 variants in the enterobacteria collection

R-GNOSIS genomes were assembled as in ^23^. pOXA-48_K8 (MT441554) was used as reference in variant calling using Snippy v4.6.0 and plasmids sharing 72% of the pOXA-48 core-genome were selected (n=224). Then, nucleotide variants in 48,500-48,853 and 14,883-16,638 zones were discarded. Mutations in 48,500-48,853 were discarded as they were identified by Sanger sequencing (Macrogen, Spain) as false positives during assembly (Illumina data). This zone contains highly repeated nucleotides which Illumina cannot resolve properly. In 14,883-16,638 pOXA-48 contained a group-II intron (*ltrA*). *ltrA* sequence was blasted (BLASTn^35^ v2.11.0) against the assemblies of each strain to confirm its presence/absence in each PVs. Identity differences in *ltrA* sequence were not considered for PVs. Insertions were not detected in hybrid assemblies and were not considered for strains sequenced just with Illumina technology. However, we could manually detect a *bla*_CTX-M-15_ gene insertion in position 7,018 in PV-D, by comparing assemblies and *in vitro* validating by PCR. Deletions between PVs were *in silico* detected with BRIG v0.95^36^ and validated by PCR amplification. We defined PVs as pOXA-48-like plasmids isolated in R-GNOSIS that share at least a 72% core-genome with pOXA-48_K8 but presented SNPs and/or indels when compared to it.

### Introducing pOXA-48 variants into bacterial isolates

A subset of 14 PVs was selected for further investigation based on the following criteria: i) PVs carrying non-synonymous mutations/deletions covering a wide representation of different genes and functions and avoiding PVs with redundant mutations in the same genes, ii) PVs carrying insertions and large rearrangements and iii) PVs with intergenic mutations near to housekeeping plasmid genes, such as genes involved in replication, conjugation or partition. Wild-type strains (donors) and *E. coli* J53 (recipient) were streaked from freezer stocks onto solid LB agar with antibiotic selection: ertapenem (0.5 mg/L) and sodium azide (100 mg/L), respectively and incubated overnight at 37°C. Several donor colonies and one recipient colony were independently inoculated in 2 mL of LB in 15-mL tubes and cultured for 6 hours (37°C and 250 rpm, Thermo Scientific™ MaxQ™ 8000). Cultures were centrifuged (15 minutes, 1,500 g) and cells were mixed in 1:2 proportion (donor:recipient) and spotted onto solid LB medium overnight at 37ºC. Transconjugants were selected by streaking the mix on LB with ertapenem and sodium azide. The presence of PVs in bacteria was confirmed by Illumina sequencing (MIGS). Additionally, each PVs was validated by PCR amplification and Sanger sequencing. For PV-A, which does not confer AMR, donors and recipients were mixed in 10:1 proportion and transconjugants were selected with sodium azide for *E. coli* J53 or Streptomycin 100 mg/L for the wild-type strain. The presence of the plasmid was confirmed by PCR screening multiple colonies. PV-B was isolated with the NucleoBond Xtra Midi Plus kit (MACHEREY-NAGEL, USA), and introduced into bacteria by electroporation as in ^37^. Transformants were selected in LB agar with amoxicillin 200 mg/L + clavulanic acid 40mg/L.

### De novo assembling and genomic analysis of E. coli J53 carrying different PVs

Genomes were assembled using SPAdes^38^ v3.15.2. Assembly quality was assessed with Quast ^39^ v5.0.2. All assemblies reached a size of 4.6-4.8 Mb and contigs >500 bp count was under 110. Prokka^40^ v1.14.6 was used to annotate genomes. Snippy v4.6.0 (https://github.com/tseemann/snippy) was used to identify variants in the *E. coli* J53 genome by mapping Illumina reads back to its assembly. Variants in the J53 strains carrying PVs were called with Snippy and breseq^41^ v0.35.6 using the annotated *E. coli* J53 genome as reference. Variants matching in J53 were discarded as assembly errors. From breseq output only predicted mutations and unassigned missing coverage (MC) were analysed because of Illumina data limitations. Snippy was used in a reverse approach, mapping the reads of *E. coli* J53 against the assemblies of the PVs-carriers. Unidentified mutations not identified in both comparisons and by both software were discarded. For pOXA-48 analysis Snippy and breseq were run using as reference pOXA-48_K8 (MT441554). Only mutations called by both programs, as well as MC and JC from breseq, were considered. The sequence of the *ltrA* gene was blasted (BLASTn^35^ v2.11.0) against the assemblies of J53 and the PVs carriers. The contig containing *ltrA* had similar length in all assemblies and different coverage than of chromosomal contigs, indicating that the *ltrA* gene did not move into the chromosome of J53. Plasmid replicons were detected with ABRicate v1.0.1 (https://github.com/tseemann/abricate) using the *plasmidfinder* database^42^. *Resfinder* database^43^ and ABRicate were used to discard the presence of other resistance genes (Supp. Table 2).

### High throughput relative fitness determination by competition assays

Competition assays were performed by using GFP-tagged strains to distinguish between populations with flow cytometry (CytoFLEX Platform Beckman Coulter Life Sciences, USA). Parameters were: 50 μl min−1 flow rate, 22 μm core size, and 10,000 events per well. Competitions were performed by competing each genotype against a GFP-tagged strain. In *E. coli* J53, each genotype had 6 replicates and the common competitor was the plasmid-free strain with pBGC^24^. In clinical strains, competitions were performed by competing each genotype (plasmid-free, and the same strain carrying different PVs against a common competitor). Each genotype was obtained from independent conjugation/transformation events, resulting in 3 independent replicates measured 6 times each (n=18 for each genotype). The common competitor for clinical strains were HKH*/PV-K + pBGA, JWC*/PV-A + pBGC and WDV*/PV-B + pBGA for each case. The common competitors carried PVs to avoid conjugative transfer during competition assays through plasmid exclusion mechanisms^44^.

Note that pBGA only differs from pBGC in the AMR gene (apramycin & chloramphenicol resistance respectively). These plasmids contain a *gfp* gene which is under the control of the P_BAD_ promoter, so GFP production is controlled by the presence of L-Arabinose. Pre-cultures were incubated overnight in LB in 96-well plates at 250 rpm (Thermo Scientific™ MaxQ™ 8000) 37ºC, then mixed 1:1 and diluted 400-fold in 200 µl of fresh LB in 96-well plates (Thermo Scientific, Denmark), and incubated during 24 in the same conditions. The initial populations were mixed (1:1) followed by diluting 400-fold in 200 µl of NaCl 0.9% with L-arabinose 0.5 % (Sigma, Spain) and incubated at 37 ºC at 250 rpm during 1.5 hours to induce GFP expression. After 24 hours, final proportions were determined as described above. The fitness of each strain relative to the GFP-tagged one was determined using equation (1):

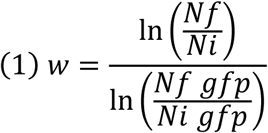

Where *w* is the relative fitness of each strain carrying a determined pOXA-48 variant compared to the GFP-tagged competitor. Ni and Nf are the number of cells of gfp-free clones at the beginning (Ni) and end (Nf) of the competition. Ni gfp and Nf gfp are the number of cells of the common GFP-tagged competitor at the beginning and end of the competition respectively. We discarded PVs loss during the competition by growing individually plasmid-carrying bacteria on LB agar and plasmid-selective antibiotics and counting colony forming units per mL at the beginning and end of the assay (PV-A was tested by PCR). Relative fitness (*w*) was normalised using the *w* from the common competitors in each case. An underrepresentation of plasmid costs during the competition assay in *E. coli* J53 due to conjugative transfer was also discarded by comparing growth curve data (using area under the growth curve, AUC) with relative fitness (Supp. Fig. 2).

### Growth curves

Growth curves were performed as in ^45^. Briefly, strains were streaked from freezer stocks onto solid LB-agar and incubated overnight at 37°C. The next day single colonies were grown in 2 mL of LB and incubated overnight at 37°C with continuous shaking (250 rpm, Thermo Scientific™ MaxQ™ 8000). Six overnight cultures were diluted 1:1,000 into fresh LB in flat-bottom 96-well plates (Thermo Scientific, Denmark), which were incubated during 24 hours at 37 ºC 250 rpm. Optical densities (OD600) were measured every 10 minutes during the incubation in a plate reader (Synergy HTX Multi-Mode Reader, BioTek Instruments, USA). The area under the growth curve (AUC) was determined by using the *growthrates* v0.8.2 & *flux* v0.3-0 packages in *Rstudio* 2021.09.2+382. When determining plasmid-variants cost in *E. coli* J53, normalised AUC was calculated by dividing the AUC of each pOXA-48-carrying isolate by the average value of the AUC of the pOXA-48-free isolate from each plate.

### Antimicrobial susceptibility testing

Bacterial AMR profile was determined by (i) LB-growth curves in the presence of different antibiotics and (ii) calculation of inhibitory concentration 90 (IC90) which corresponds to the antibiotic concentration inhibiting 90% of the bacterial growth in the absence of antibiotics. For (i) we used the protocol described above and for (ii) strains were streaked from freezer stocks onto solid MH-agar medium and incubated overnight (37ºC). Then, single colonies of bacterial cells (n=5 or 10) were inoculated in parallel in liquid MH starter cultures and incubated at 37°C for 24 hours at 250 rpm. Later, each culture was diluted 1:1,000 in MH medium (∼10^6^ cfu) and 200 µl of the final solution were added to a flat-bottom 96-well plate (Thermo Scientific, Denmark) containing the appropriate antibiotic concentration. Antibiotics tested were ertapenem, meropenem and amoxicillin+clavulanic acid. IC90 values were measured after 24 hours of incubation (37ºC). Optical density at 600 nm (OD600) was determined in a Synergy HTX (BioTek Instruments, USA) plate reader after 30 seconds of orbital shaking. MH containing each antibiotic concentration was used as blank.

### Determining plasmid transfer rate in E. coli J53

PVs transfer rate was evaluated using *E. coli* J53 as donor and *E. coli* J53/pBGC (a non-mobilizable and chloramphenicol-resistant plasmid) as recipient. Donors and recipients were streaked in selective agar (ertapenem 0.5mg/L or chloramphenicol 50µg/mL, respectively). After an overnight incubation at 37ºC, colonies of each donor and the recipient strain were independently inoculated in 2 mL of LB in 15-mL culture tubes and incubated overnight at 37 °C and 250 rpm (Thermo Scientific™ MaxQ™ 8000). Then, 100 µl of donor and recipient were mixed in a 1:1 proportion and incubated on a LB agar plate at 37°C for 2 hours. Subsequently, serial dilutions of each mix were prepared in sterile NaCl 0.9% and plated on selective media for each genotype (carbenicillin 100 µg mL^−1^, Chloramphenicol 50µg/mL and both antibiotics together). Conjugation rates were determined using the end-point method for solid surfaces as in ^23^.

### Plasmid copy number determination by quantitative PCR (qPCR)

Each genotype was streaked in LB agar and incubated overnight at 37ºC. The next day 2 independent colonies were resuspended in 800 µl in sterile water (Fisher Scientific, Spain) and boiled for 10 minutes (95ºC). Each sample was centrifuged to spin down cellular debris. Then, 3 independent reactions per colony were performed in triplicate, with 1 µl of the supernatant as DNA template and using with the NZYSupreme qPCR Green Master Mix (2x), ROX plus kit (NZYtech, Portugal) and the 7500 Real Time PCR System (Applied Biosystems, USA). Targeted plasmid and chromosome genes were *bla*_OXA-48_ (amplicon size 200 pb; efficiency 97.35-98.09%, r^2^=0.996-0.986) & *dnaE* (chromosomal gene with one copy, amplicon size 200 bp; efficiency 98.44-100.64%, r^2^= 0.989-0.996) respectively. The efficiency was calculated using serial ¼ dilutions of K8 strain (PV-I) and J53/PV-I as in ^46^. The amplification conditions were: 5 minutes denaturation (95ºC) followed by 30 cycles of 15 seconds denaturation, 30 seconds annealing (55ºC) and 30 seconds extension (60ºC). The relative plasmid copy number was calculated using equation (2):

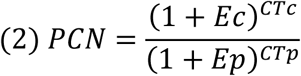

where PCN is the plasmid copy number per chromosome, Ec and Ep are the efficiencies of the chromosomal and plasmid reactions (relative to 1), and CTc and CTp are the threshold cycles for chromosomal and plasmid reactions.

### Curing pOXA-48 like plasmids using a CRISPR/Cas9-based system and reintroducing PVs into the clinical isolates

Two different plasmid versions carrying CRISPR/Cas9 machinery were used in this project: pLC10-Kan (kanamycin resistant) and pLC10-Apra (apramycin resistant). pLC10-Kan was used in JWC* and HKH* and pLC10-Apra in WDV*. First pOXA-48 carrying strains were made competent following the protocol described in ^36^. Then each pLC10 was introduced in the cells by electroporation using 0.1 cm cuvettes and 1.8 kV pulse (MicroPulser Electroporator, Biorad Spain). Transformants were selected on LB agar plates with kanamycin 250- 512mg/L or apramycin 30mg/L for each case. Transformants were verified by PCR (Supp. Table 4). Then, CRISPR/Cas9 machinery was induced by resuspending several transformant colonies in 500 µl of LB with kanamycin or apramycin, 0.2 mg/L anhydrotetracycline (aTc), to activate *Cas9* expression, and IPTG 0.08 mM to enhance sgRNA expression. Then, suspensions were incubated for 2 hours at 30ºC with agitation (250 rpm, Thermo Scientific™ MaxQ™ 8000) and was streaked and incubated overnight at 37ºC on LB agar to cure pLC10. Note that pLC10 *oriC* is based on pSC101 and codes for a thermosensitive replication protein. The next day single colonies were streaked parallelly in LB agar and LB agar supplemented with ertapenem 0.5 mg/L, kanamycin or apramycin. Only colonies that were sensitive to both antibiotics were recovered and sequenced by Illumina (MIGS, USA). Then different plasmid variants were re-introduced in triplicate (in parallel) to plasmid-free cells as described above.

### Analysis of wild-type strains involved in within-patient pOXA-48 evolution

4 potential cases of within-patient pOXA-48 evolution were identified: HKH, JWC, WDV and HAX. HAX was discarded because the PVs differed from each other just by a synonymous SNP (Supp. Table 1). Unicycler^47^ v0.4.9 with default parameters was used to obtain hybrid-assemblies from K153, K229, C288, C289, K163 and K165 strains. Long-reads were also assembled with Flye^48^ v2.9 and circularization was confirmed in Bandage^49^ v0.8.1. Medaka v1.4.3 (https://github.com/nanoporetech/medaka) was used to obtain consensus sequences. Several rounds of Pilon^50^ v1.24 were performed mapping the trimmed Illumina reads. Contigs were rotated with Circlator fixstart^51^ v1.5.5. Long-read assembly quality was controlled in IGV^52^ v2.11.1. PVs assemblies were confirmed by mapping short- and long-reads and by aligning the assemblies to the reference pOXA-48_K8 (MT441554) with BWA-MEM^53^ v0.7.17 and minimap2^54^ v2.21. Alignments were visualized in IGV. Closed assemblies were annotated with PGAP^55^ v2021-07-01.build5508. Breseq v0.36.0 was used to identify SNPs and structural variants. To discard false-positive calls different combinations of breseq runs were performed. For K164, K165-2 cured, K165-1, K165-3, K165-4, K165-5, K165-6, K165-7 and K166 (JWC), K151, K152, K229 cured (WDV) and C289 cured (HKH), the trimmed reads were mapped to the closed strains from their respective patients. ABRicate v1.0.1 with the *plasmidfinder* and *resfinder* databases was used to confirm clonality and isogeneity between within-patient evolved and cured strains. Further details on workflow and analysis criteria are provided in https://github.com/LaboraTORIbio/within_patient_evolution.

### Construction of phylogenetic trees

Snippy v4.6.0 was used to find SNPs between all *E. coli* ST10 & ST744 (reference C288), *K. pneumoniae* ST11 (reference K153) and K. pneumoniae ST307 (reference K163) from the R-GNOSIS collection. Strain K25 (ST11) was removed from the analysis because the fastq files were truncated. Snippy-core (https://github.com/tseemann/snippy) was used to find the core genome. Strain K78 (ST11) was removed for diverging too much from the rest of the strains. Gubbins^56^ v3.1.4 was used to remove recombinant regions and SNPs were extracted with snp-sites^57^ v2.5.1. Maximum-likelihood trees were constructed with IQ-TREE^58^ v1.6.12 from the extracted alignments with best evolutionary model detection and an ultrafast bootstrap of 1000 optimized by hill-climbing nearest neighbour interchange (NNI) on the corresponding bootstrap alignment. Trees were visualized and edited in iTOL^59^ and Inkscape v0.17.

### Statistical analyses

All statistical analyses were performed in *Rstudio* 2021.09.2+382 (R v4.1.1 2021- 08-10) with packages *rstatix* v0.7.0, *tidyverse* v1.3.1 and *car* v3.0-12. To test homoscedasticity and the normality of data for each dataset, Shapiro-Wilk test, Levene’s Test and Bartlett’s test were performed. Then according to each data structure parametric and nonparametric tests were performed (see main manuscript for each test).

## Acknowledgments

We appreciate the technical support of Laura Jaraba Soto. We also thank Craig MacLean, José Penadés, José Antonio Escudero and Daniel Padfield for constructive comments. This work was supported by the European Research Council under the European Union’s Horizon 2020 research and innovation programme (ERC grant agreement no. 757440-PLASREVOLUTION) and by the Instituto de Salud Carlos III (PI19/00749) co-funded by European Development Regional Fund ‘a way to achieve Europe’. The R-GNOSIS project received financial support from the European Commission (grant no. R-GNOSIS-FP7- HEALTH-F3-2011-282512). A.S.-L. is supported by the European Commission (H2020-MSCA-IF-2019, 895671-REPLAY) and by the European Society of Clinical Microbiology and Infectious Diseases (ESCMID, Research Grant 2022). J.R.-B. acknowledges financial support by a Miguel Servet contract from Instituto de Salud Carlos III (ISCIII) (grant no. CP20/00154), co-funded by ESF, ‘Investing in your future’, CIBERINFEC, co-funded with FEDER funds, and project PI21/01363, funded by Instituto de Salud Carlos III (ISCIII) and co-funded by the European Union.

## Author contributions

A.S.M and J.DF were responsible for the conceptualization of the study; J.DF, L.C, D.B. and A.S.M designed the methodology. L.T.-C, J.DF and R.L.-S analysed the genomic data; C.C, A.S.-L., A.A.V and J.DF performed experiments and contributed to data analysis; R.C. designed and supervised sampling and collection of bacterial isolates. M.H.-G. collected the bacterial isolates. J.DF and A.S.M analysed data and prepared the original draft of the manuscript and undertook the reviewing and editing; All authors supervised and approved the final version of the manuscript; A.S.M was responsible for funding acquisition and supervision.

## Data availability

The sequence data that support the findings of this study are available in the National Center for Biotechnology Information Database with the accession code PRJNA838107 (https://www.ncbi.nlm.nih.gov/bioproject/838107). The remaining R-GNOSIS sequences can be found in ^23^.

## Supplementary figures

**supp. Fig. 1.**
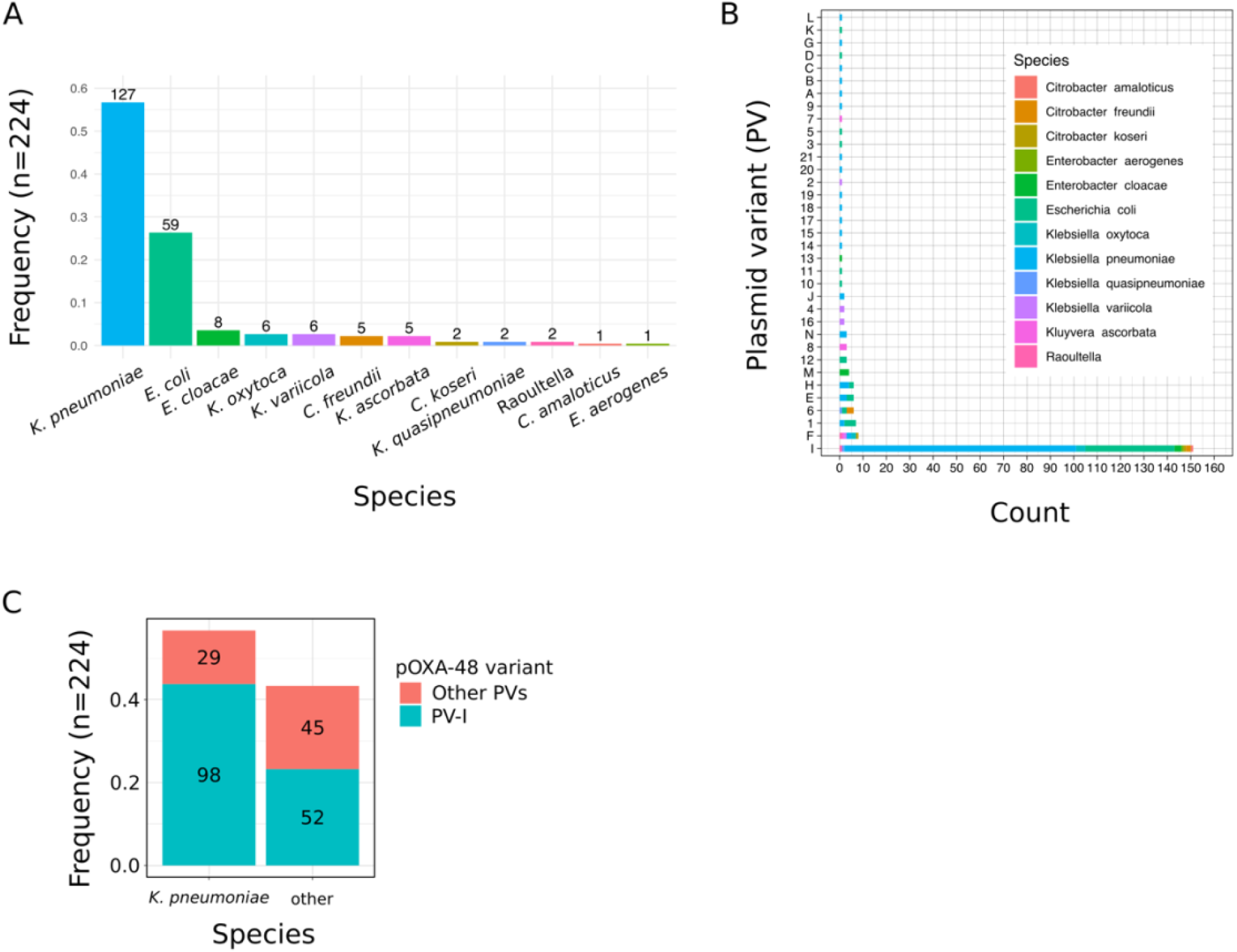
Enterobacteria carrying pOXA-48 recovered during the R-GNOSIS study. A) Frequency of clinical isolates by species. Numbers on top of the bars indicate the number of isolates. B) Distribution of PVs from the collection by count and species (colours). C) Frequency of isolates of *K. pneumoniae* or other enterobacteria carrying the most common pOXA-48 variant, PV-I. Colours correspond to the PVs variant and the number within the bars correspond to the isolate count.

**supp. Fig. 2.**
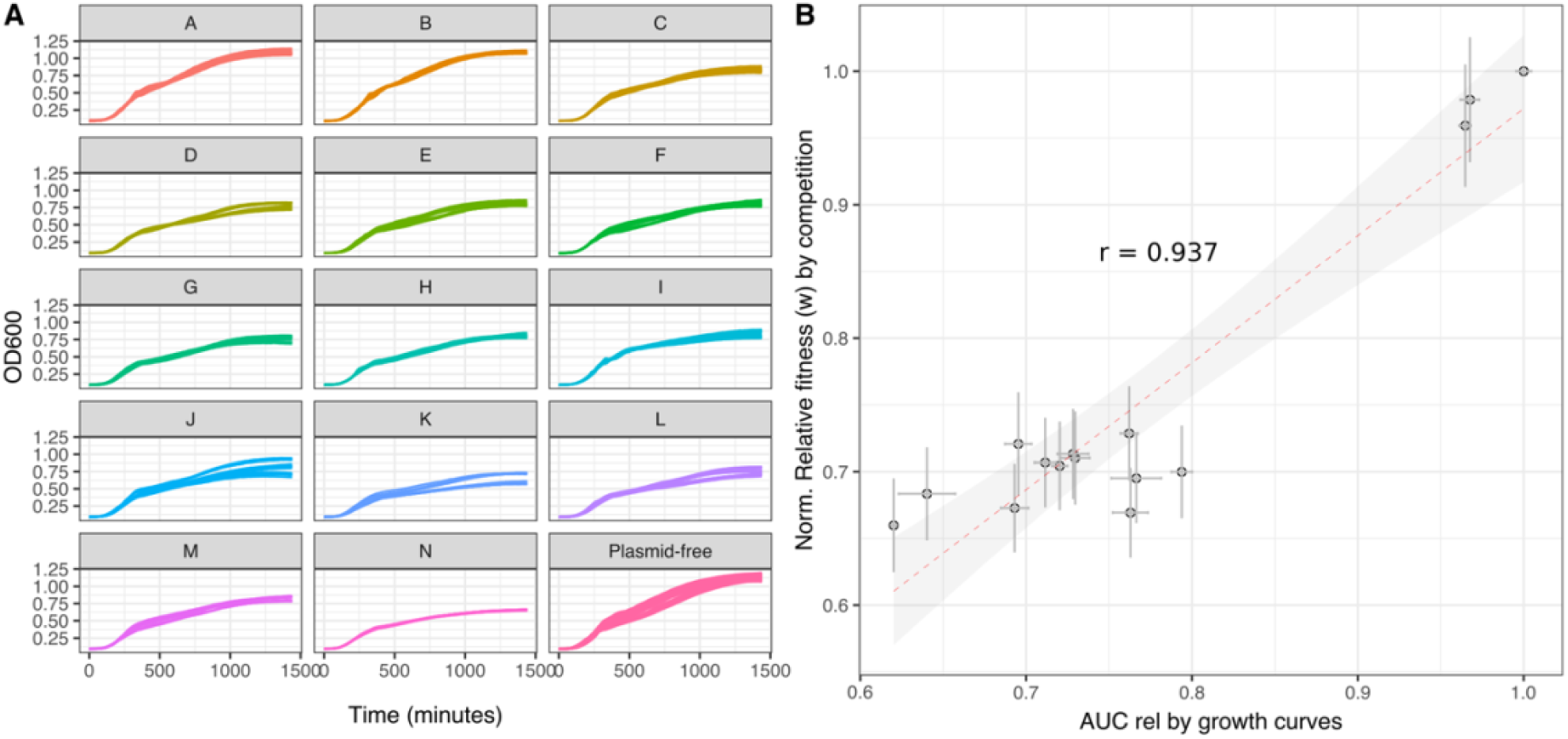
Growth dynamics of *E. coli* J53 carrying different PVs. A) Growth curves of *E. coli* J53 carrying different PVs. Vertical axis shows the optical density at 600 nm (OD600) and the horizontal axis time in minutes. Each PV is indicated in the top label. B) Linear correlation of relative fitness (*w*) calculated by competition assays or by area under the growth curves (Pearson’s product-moment correlation t = 9.6665, df = 13, P = 2.663e-07, cor 0.9369456). Lines indicate the propagated standard error of the mean and points indicate the mean values for each genotype.

**supp. Fig. 3.**
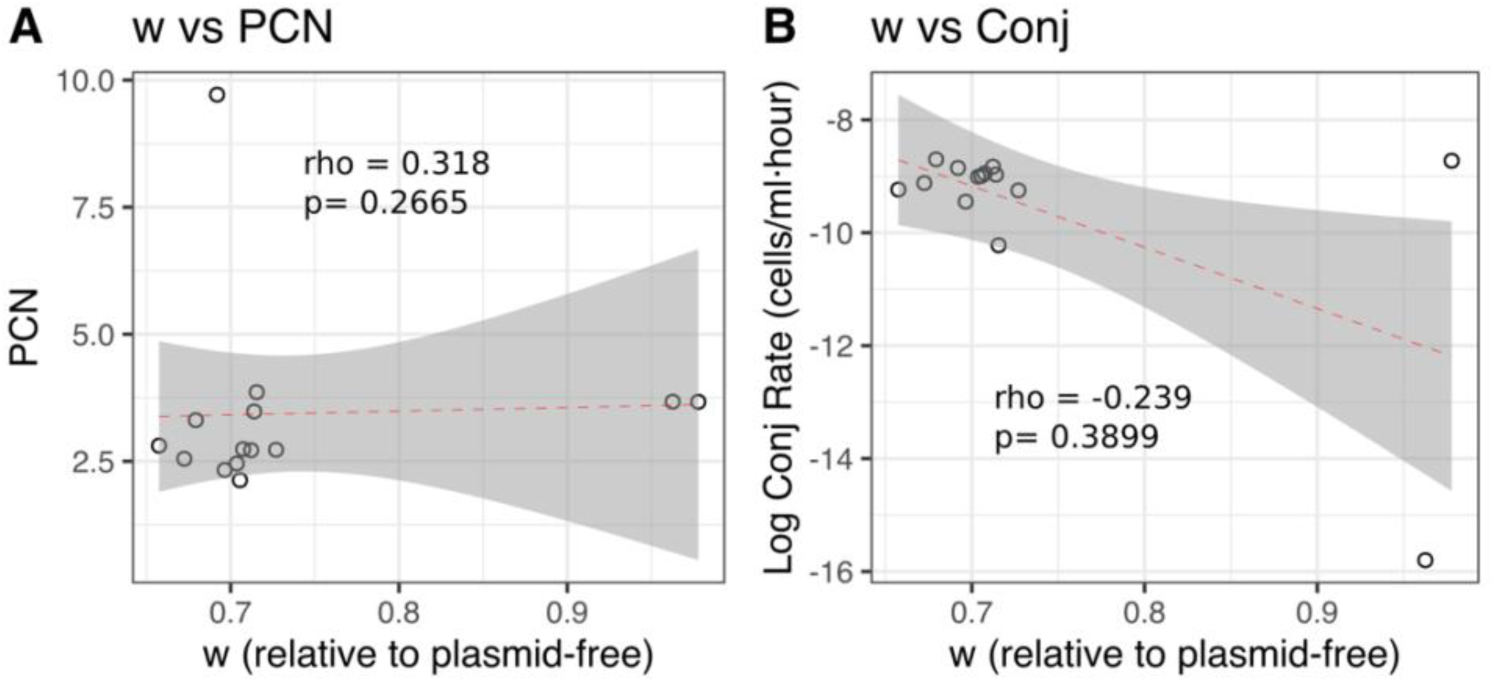
Plasmid copy number (PCN) and conjugation rates do not correlate with plasmid fitness costs. Correlation between relative fitness (*w*) and A) PCN, or B) log_10_ conjugation rate of *E. coli* J53 carrying different PVs relative to the plasmid-free strain. In each panel individual dots correspond to the median values of of *E. coli* J53 carrying different PVs. Spearman’s rank correlation rho and p-value (p) for each case are indicated in the figure.

**supp. Fig. 4.**
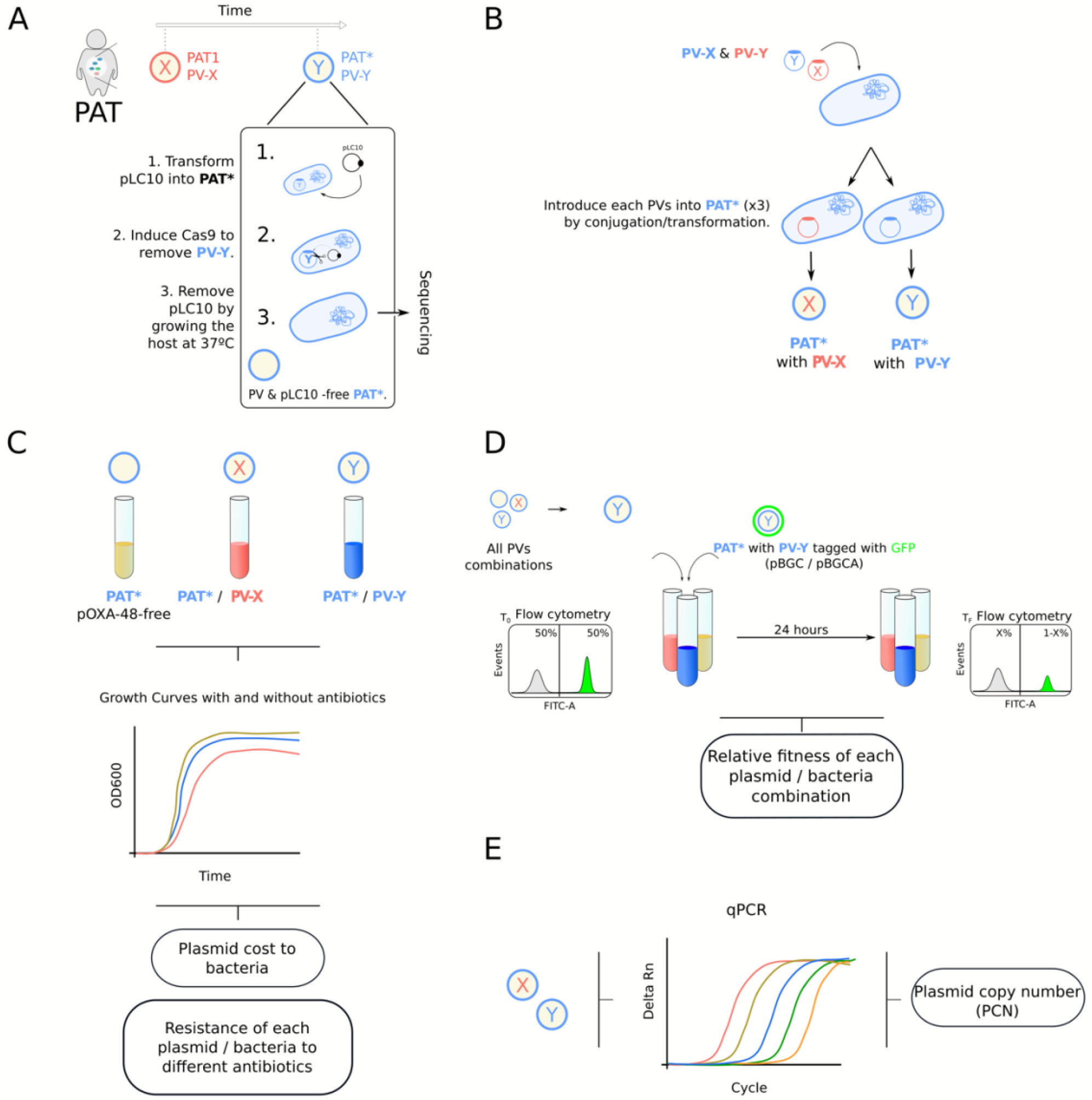
Workflow used to explore within-patient AMR evolution. A) PVs curing from clinical isolates; B) re-introduction of different PVs into the clinical isolates; C) evaluation of the plasmid-cost and the resistance profile of each plasmid-carrying bacteria combination; D) relative fitness (*w*) calculation; and E) calculation of plasmid copy number (PCN) for each PV.

**supp. Fig. 5.**
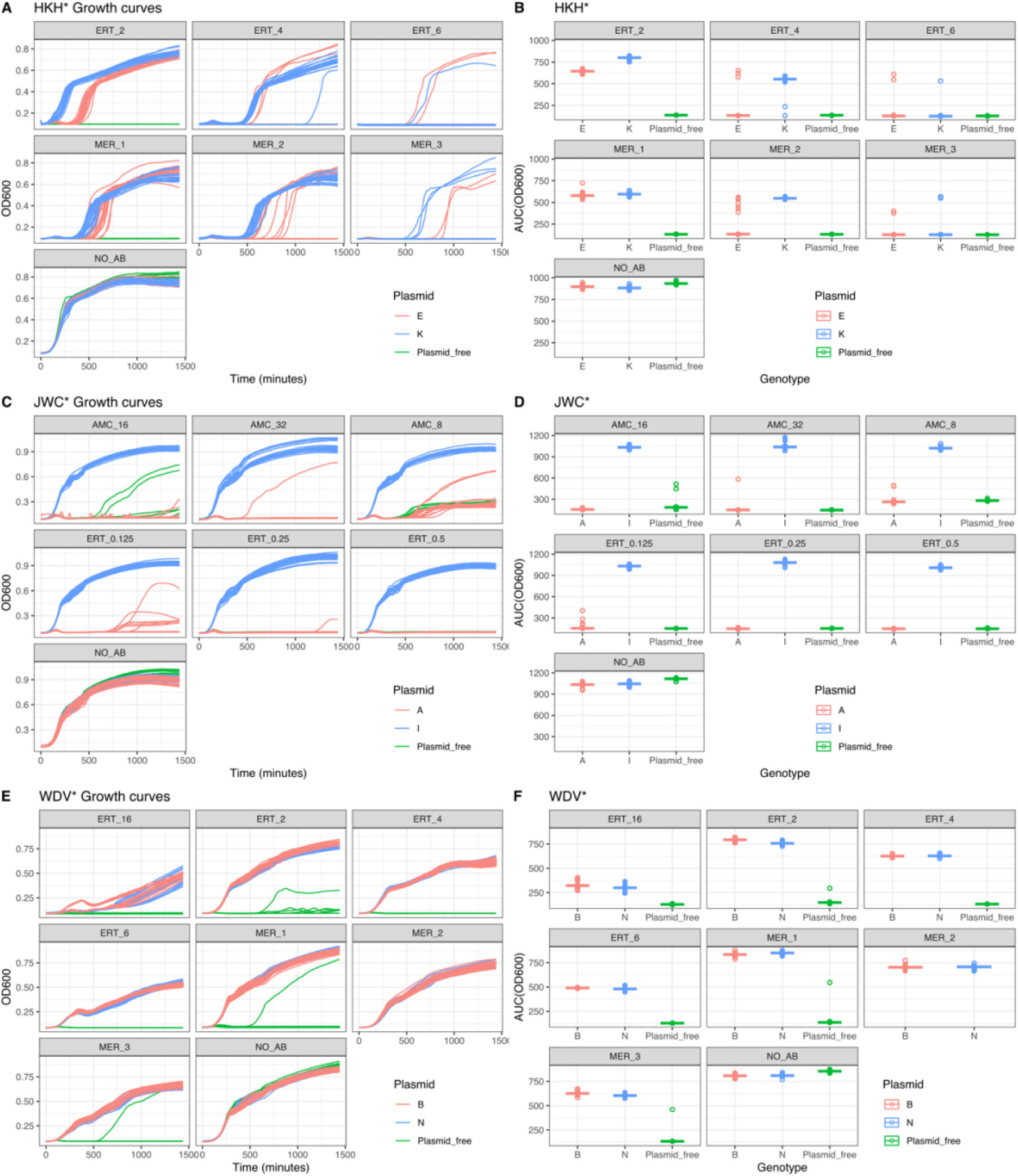
Growth dynamics of the clinical bacteria carrying different PVs isolated from the three patients under different antibiotic treatments. A) Growth curves of HKH* carrying different PVs (indicated by different colours, see legend). Vertical axis shows the OD600 and horizontal axis the time in minutes. Each antibiotic concentration is indicated in the top label (ERT stands for ertapenem; MER for meropenem and NO_AB for no antibiotic treatment, the number indicates the concentration in mg/L). n=18 for each genotype and treatment. B) Growth of different HKH* carrying different PVs (as in A), using the values of the area under the curve (AUC in vertical axis, t = 1500 minutes). Individual points indicate individual values (n=18 for each genotype and treatment) and horizontal lines indicate the median value of the replicates. C) Growth curves of JWC* carrying different PVs (as in A). AMC stands for amoxicillin + clavulanic acid. D) Growth of different JWC* carrying different PVs using the values of the area under the curve (as in B). E) Growth curves of WDV* carrying different PVs (as in A and C). F) Growth of different WDV* carrying different PVs using the values of the area under the curve (as in B and D).

## References

1. Murray, C. J. et al. Global burden of bacterial antimicrobial resistance in 2019: a systematic analysis. The Lancet 399, 629–655 (2022).

2. Vincent, J.-L. et al. International study of the prevalence and outcomes of infection in intensive care units. JAMA 302, 2323–9 (2009).

3. van Schaik, W. The human gut resistome. Philosophical Transactions of the Royal Society B: Biological Sciences 370, 20140087 (2015).

4. Partridge, S. R., Kwong, S. M., Firth, N. & Jensen, S. O. Mobile Genetic Elements Associated with Antimicrobial Resistance. Clinical Microbiology Reviews 31, (2018).

5. Dimitriu, T., Matthews, A. C. & Buckling, A. Increased copy number couples the evolution of plasmid horizontal transmission and plasmid-encoded antibiotic resistance. Proceedings of the National Academy of Sciences 118, e2107818118 (2021).

6. San Millan, A., Escudero, J. A., Gifford, D. R., Mazel, D. & MacLean, R. C. Multicopy plasmids potentiate the evolution of antibiotic resistance in bacteria. Nature Ecology & Evolution 1, 0010 (2017).

7. Wheatley, R. et al. Rapid evolution and host immunity drive the rise and fall of carbapenem resistance during an acute Pseudomonas aeruginosa infection. doi:10.1038/s41467-021-22814-9.

8. Fröhlich, C. et al. Cryptic β-Lactamase Evolution Is Driven by Low β-Lactam Concentrations. mSphere 6, (2021).

9. Souque, C., Escudero, J. A. & MacLean, R. C. Integron activity accelerates the evolution of antibiotic resistance. Elife 10, (2021).

10. Bottery, M. J., Wood, A. J. & Brockhurst, M. A. Adaptive modulation of antibiotic resistance through intragenomic coevolution. Nature Ecology & Evolution 1, 1364–1369 (2017).

11. Vogwill, T. & MacLean, R. C. The genetic basis of the fitness costs of antimicrobial resistance: a meta-analysis approach. Evolutionary Applications 8, 284–295 (2015).

12. Martínez-García, L., González-Alba, J. M., Baquero, F., Cantón, R. & Galán, J. C. Ceftazidime Is the Key Diversification and Selection Driver of VIM-Type Carbapenemases. mBio 9, (2018).

13. Brockhurst, M. A. & Harrison, E. Ecological and evolutionary solutions to the plasmid paradox. Trends in Microbiology (2021) doi:10.1016/j.tim.2021.11.001.

14. Loftie-Eaton, W. et al. Compensatory mutations improve general permissiveness to antibiotic resistance plasmids. Nature Ecology & Evolution 1, 1354–1363 (2017).

15. Hall, J. P. J. et al. Plasmid fitness costs are caused by specific genetic conflicts enabling resolution by compensatory mutation. PLoS Biology 19, (2021).

16. Rajer, F. & Sandegren, L. The Role of Antibiotic Resistance Genes in the Fitness Cost of Multiresistance Plasmids. mBio 13, e0355221 (2022).

17. Humphrey, B. et al. Fitness of Escherichia coli strains carrying expressed and partially silent IncN and IncP1 plasmids. BMC Microbiology 12, 1–9 (2012).

18. Andersson, D. I. & Hughes, D. Antibiotic resistance and its cost: Is it possible to reverse resistance? Nature Reviews Microbiology vol. 8 260–271 (2010).

19. Basra, P. et al. Fitness Tradeoffs of Antibiotic Resistance in Extraintestinal Pathogenic Escherichia coli. Genome Biology and Evolution 10, 667–679 (2018).

20. Bonomo, R. A. et al. Carbapenemase-Producing Organisms: A Global Scourge. Clinical Infectious Diseases 66, 1290–1297 (2018).

21. David, S. et al. Epidemic of carbapenem-resistant Klebsiella pneumoniae in Europe is driven by nosocomial spread. Nature Microbiology 4, 1919–1929 (2019).

22. Cassini, A. et al. Attributable deaths and disability-adjusted life-years caused by infections with antibiotic-resistant bacteria in the EU and the European Economic Area in 2015: a population-level modelling analysis. The Lancet Infectious Diseases 19, 56–66 (2019).

23. León-Sampedro, R. et al. Pervasive transmission of a carbapenem resistance plasmid in the gut microbiota of hospitalized patients. Nature Microbiology 6, 606–616 (2021).

24. Alonso-del Valle, A. et al. Variability of plasmid fitness effects contributes to plasmid persistence in bacterial communities. Nature Communications 12, 2653 (2021).

25. Hernández-García, M. et al. Characterization of carbapenemase-producing Enterobacteriaceae from colonized patients in a university hospital in Madrid, Spain, during the R-GNOSIS project depicts increased clonal diversity over time with maintenance of high-risk clones. Journal of Antimicrobial Chemotherapy 73, 3039–3043 (2018).

26. Matsumura, Y., Peirano, G. & Pitout, J. D. D. Complete Genome Sequence of Escherichia coli J53, an Azide-Resistant Laboratory Strain Used for Conjugation Experiments. Genome Announcements 6, (2018).

27. Yurtsev, E. A., Chao, H. X., Datta, M. S., Artemova, T. & Gore, J. Bacterial cheating drives the population dynamics of cooperative antibiotic resistance plasmids. Molecular Systems Biology 9, (2013).

28. zur Wiesch, P. A., Kouyos, R., Engelstädter, J., Regoes, R. R. & Bonhoeffer, S. Population biological principles of drug-resistance evolution in infectious diseases. The Lancet Infectious Diseases 11, 236–247 (2011).

29. Caballero, J. D. et al. Polyclonal pathogen populations accelerate the evolution of antibiotic resistance in patients. (2022) doi:10.1101/2021.12.10.472119.

30. Stracy, M. et al. Minimizing treatment-induced emergence of antibiotic resistance in bacterial infections. Science (1979) 375, 889–894 (2022).

31. Yang, J. et al. High-Efficiency Scarless Genetic Modification in Escherichia coli by Using Lambda Red Recombination and I-SceI Cleavage. Applied and Environmental Microbiology 80, 3826–3834 (2014).

32. Goldberg, G. W., Jiang, W., Bikard, D. & Marraffini, L. A. Conditional tolerance of temperate phages via transcription-dependent CRISPR-Cas targeting. Nature 514, 633 (2014).

33. Jiang, Y. et al. Multigene Editing in the Escherichia coli Genome via the CRISPR-Cas9 System. (2015) doi:10.1128/AEM.04023-14.

34. Engler, C., Kandzia, R. & Marillonnet, S. A One Pot, One Step, Precision Cloning Method with High Throughput Capability. PLOS ONE 3, e3647 (2008).

35. Altschul, S. F., Gish, W., Miller, W., Myers, E. W. & Lipman, D. J. Basic local alignment search tool. Journal of Molecular Biology 215, 403–410 (1990).

36. Alikhan, N. F., Petty, N. K., ben Zakour, N. L. & Beatson, S. A. BLAST Ring Image Generator (BRIG): Simple prokaryote genome comparisons. BMC Genomics 12, 1–10 (2011).

37. Fournet-Fayard, S., Joly, B. & Forestier, C. Transformation of wild type Klebsiella pneumoniae with plasmid DNA by electroporation. Journal of Microbiological Methods 24, 49–54 (1995).

38. Prjibelski, A., Antipov, D., Meleshko, D., Lapidus, A. & Korobeynikov, A. Using SPAdes De Novo Assembler. Current Protocols in Bioinformatics 70, (2020).

39. Mikheenko, A., Prjibelski, A., Saveliev, V., Antipov, D. & Gurevich, A. Versatile genome assembly evaluation with QUAST-LG. Bioinformatics 34, i142–i150 (2018).

40. Seemann, T. Prokka: rapid prokaryotic genome annotation. Bioinformatics 30, 2068–2069 (2014).

41. Deatherage, D. E. & Barrick, J. E. Identification of mutations in laboratory evolved microbes from next-generation sequencing data using breseq. Methods Mol Biol 1151, 165 (2014).

42. Carattoli, A. et al. In Silico detection and typing of plasmids using plasmidfinder and plasmid multilocus sequence typing. Antimicrobial Agents and Chemotherapy 58, 3895–3903 (2014).

43. Zankari, E. et al. Identification of acquired antimicrobial resistance genes. Journal of Antimicrobial Chemotherapy 67, 2640–2644 (2012).

44. Garcillán-Barcia, M. P. & de la Cruz, F. Why is entry exclusion an essential feature of conjugative plasmids? Plasmid 60, 1–18 (2008).

45. DelaFuente, J., Rodriguez-Beltran, J. & San Millan, A. Methods to Study Fitness and Compensatory Adaptation in Plasmid-Carrying Bacteria. Methods in Molecular Biology 2075, 371–382 (2020).

46. Millan, A. S. et al. Small-plasmid-mediated antibiotic resistance is enhanced by increases in plasmid copy number and bacterial fitness. Antimicrob Agents Chemother 59, 3335–3341 (2015).

47. Wick, R. R., Judd, L. M., Gorrie, C. L. & Holt, K. E. Unicycler: Resolving bacterial genome assemblies from short and long sequencing reads. PLOS Computational Biology 13, e1005595 (2017).

48. Kolmogorov, M., Yuan, J., Lin, Y. & Pevzner, P. A. Assembly of long, error-prone reads using repeat graphs. Nature Biotechnology 2019 37:5 37, 540–546 (2019).

49. Wick, R. R., Schultz, M. B., Zobel, J. & Holt, K. E. Bandage: interactive visualization of de novo genome assemblies. Bioinformatics 31, 3350–3352 (2015).

50. Walker, B. J. et al. Pilon: An Integrated Tool for Comprehensive Microbial Variant Detection and Genome Assembly Improvement. PLOS ONE 9, e112963 (2014).

51. Hunt, M. et al. Circlator: Automated circularization of genome assemblies using long sequencing reads. Genome Biology 16, 1–10 (2015).

52. Robinson, J. T. et al. Integrative genomics viewer. Nature Biotechnology 2011 29:1 29, 24–26 (2011).

53. Li, H. & Durbin, R. Fast and accurate long-read alignment with Burrows– Wheeler transform. Bioinformatics 26, 589–595 (2010).

54. Li, H. Minimap2: pairwise alignment for nucleotide sequences. Bioinformatics 4, 3094–3100 (2018).

55. Tatusova, T. et al. NCBI prokaryotic genome annotation pipeline. Nucleic Acids Research 44, 6614–6624 (2016).

56. Croucher, N. J. et al. Rapid phylogenetic analysis of large samples of recombinant bacterial whole genome sequences using Gubbins. Nucleic Acids Research 43, e15–e15 (2015).

57. Page, A. J. et al. SNP-sites: rapid efficient extraction of SNPs from multi-FASTA alignments. Microb Genom 2, e000056 (2016).

58. Nguyen, L. T., Schmidt, H. A., von Haeseler, A. & Minh, B. Q. IQ-TREE: A Fast and Effective Stochastic Algorithm for Estimating Maximum-Likelihood Phylogenies. Molecular Biology and Evolution 32, 268–274 (2015).

59. Letunic, I. & Bork, P. Interactive Tree Of Life (iTOL) v5: an online tool for phylogenetic tree display and annotation. Nucleic Acids Research 49, W293–W296 (2021).

